# ICOR: Improving codon optimization with recurrent neural networks

**DOI:** 10.1101/2021.11.08.467706

**Authors:** Rishab Jain, Aditya Jain, Elizabeth Mauro, Kevin LeShane, Douglas Densmore

## Abstract

**Background:** In protein sequences—as there are 61 sense codons but only 20 standard amino acids— most amino acids are encoded by more than one codon. Although such synonymous codons do not alter the encoded amino acid sequence, their selection can dramatically affect the expression of the resulting protein. Codon optimization of synthetic DNA sequences is important for heterologous expression. However, existing solutions are primarily based on choosing high-frequency codons only, neglecting the important effects of rare codons. In this paper, we propose a novel recurrent-neural-network based codon optimization tool, ICOR, that aims to learn codon usage bias on a genomic dataset of *Escherichia coli*. We compile a dataset of over 7,000 non-redundant, high-expression, robust genes which are used for deep learning. The model uses a bidirectional long short-term memory-based architecture, allowing for the sequential context of codon usage in genes to be learned. Our tool can predict synonymous codons for synthetic genes toward optimal expression in *Escherichia coli*.

**Results:** We demonstrate that sequential context achieved via RNN may yield codon selection that is more similar to the host genome. Based on computational metrics that predict protein expression, ICOR theoretically optimizes protein expression more than frequency-based approaches. ICOR is evaluated on 1,481 *Escherichia coli* genes as well as a benchmark set of 40 select DNA sequences whose heterologous expression has been previously characterized. ICOR’s performance is measured across five metrics: the Codon Adaptation Index, GC-content, negative repeat elements, negative cis-regulatory elements, and codon frequency distribution.

**Conclusions:** The results, based on in silico metrics, indicate that ICOR codon optimization is theoretically more effective in enhancing recombinant expression of proteins over other established codon optimization techniques. Our tool is provided as an open-source software package that includes the benchmark set of sequences used in this study.

## Background

Designing synthetic genes for heterologous expression is a keystone of synthetic biology[1]. The expression of recombinant proteins in a heterologous host has applications from manufacturing pharmaceuticals to vaccines. For instance, producing malaria vaccine FALVAC-1[2] involves designing synthetic plasmids, transfection into the *Escherichia coli* (*E. coli*) host factory, growing the cells, and harvesting the resulting protein[3]. *E. coli* is well-established as a host for heterologous expression[4], however, codon bias limits the use of *E. coli* as an expression platform[5]. To increase the efficiency of recombinant expression in *E. coli*, improving codon optimization is an area of particular interest.

Expression levels of synthetic genes in heterologous hosts are dependent on multiple factors including codon usage bias[5–7]. During the process of translation, complimentary tRNAs are used to read codons from an mRNA strand. In *E. coli*, the frequency of a certain codon in its genome is positively correlated with the presence of tRNAs for that codon[5, 8, 9]. Thus, choosing synonymous codons that are more frequently found in a host genome may improve heterologous expression, with 2 to 15-fold increases typically measured in a *E. coli* chassis[7]. Although synonymous codons may code for the same amino acid, they are not redundant[10, 11]. In a study by Gao et al., low expression levels of human immunodeficiency virus genes in mammalian cells were attributed to rare codon usage[12]. Other studies observe that codon bias affects gene expression and even protein folding and solubility[13–15]. Therefore, it is important to understand the underlying codon usage bias of the chassis to maximize protein expression.

Today, there are a range of FDA-approved recombinant DNA products from synthetic insulin to Hepatitis B therapeutics[16]. Codon optimization tools can be used to increase protein expression towards improving the efficiency of manufacturing such products[15]. Codon optimization techniques that are based on biological indexes replace synonymous codons with the most abundant codon found in the host organism’s genome[17]. Our review shows that many industry-standard tools employ the aforementioned strategy, causing unintended consequences for the cell, such as an imbalanced tRNA pool[10, 18]. If just one codon of a synonymous set is used throughout an entire synthetic gene, when expressed in the host, metabolic stress and translational error may be imposed[19]. Research has shown that using high-frequency codons only during codon optimization leads to incorrect protein folding and the formation of insoluble proteins[20]. Further, rare codons have been found to play an important role in protein folding[21], thereby raising the interest for understanding subpatterns of codon usage along with surrounding context in which codons are used rather than synonymous codon frequency alone.

Understanding the context by which synonymous codons are used in a gene may be essential to unlock the full evolutionary-instilled potential of a cell factory towards heterologous expression. By utilizing synonymous codons based patterns and sequential context of their prevalence in the host genome, protein expression could be increased while preventing plasmid toxicity. To best learn sequential and contextual patterns of the host, deep learning can be leveraged for its high level of abstraction on large datasets[22]. Deep learning systems show promise in bioinformatics, potentially offering improvements over non-machine learning algorithms[23].

Recurrent neural networks (RNNs) are a class of deep neural networks that can grasp temporal data, thus demonstrating utility in applications that require an understanding of sequential information[24]. For example, speech recognition models that utilize long short-term memory (LSTM) architectures[25]—a type of RNN—take advantage of the memory built into the LSTM module allowing it to interpret speech based on the surrounding context. For codon optimization, RNNs may offer improved synonymous codon selection if designed to understand underlying patterns of synonymous codon usage to inform subsequent codon prediction. By treating each amino acid as a timestep in a sequence, the RNN evaluates its prediction in the context of surrounding amino acids.

In this study, a deep learning tool, ICOR – Improving Codon Optimization with RNNs – is trained on a large, robust, non-redundant dataset of *E. coli* genomes. This “big data” approach allows our model to learn codon usage across multitudinous genes of *E. coli* and develop a model to improve codon optimization by understanding context. ICOR adopts the Bidirectional Long-Short-Term Memory (BiLSTM) architecture[26] because of its ability to preserve temporal information from both the past and future. In a gene, the BiLSTM would theoretically use surrounding synonymous codons to make a prediction.

40 benchmark genes as shown in Table S1 along with 1,481 *E. coli* genes were used for testing and data analysis, allowing us to evaluate the model in two ways: its ability to optimize genes used in past recombinant expression studies, and its ability to replicate but still optimize sequences from *E. coli*. The resultant optimized sequences from the ICOR model were compared to six approaches (original, uniform random choice (URC), background frequency choice (BFC), extended random choice (ERC), highest frequency choice (HFC), GenScript’s GenSmart[27]) as outlined in S1 Appendix. GenSmart is accepted as the industry-standard benchmark due to recognition in past studies[28], ease-of-use, and accessibility. To gauge performance of the codon optimization approaches, the Codon Adaptation Index (CAI), GC-content, codon frequency distribution (CFD), negative repeat elements, negative cis-regulatory elements, and algorithm run-time are measured on the 40 benchmark and 1,481 *E. coli* genes. These in silico metrics demonstrate the effectiveness of this approach as an algorithm. The ICOR tool is open-source and can be accessed at: https://doi.org/10.5281/zenodo.5529209.

## Implementation

### Model Training Dataset

We use the National Center for Biotechnology Information’s (NCBI) GenBank database[29] which includes 6,877,000 genes reported for many *E. coli* strains. *E. coli* genes that were shorter than 90 amino acids in length were removed due to their hypothetical or specialized nature. Then, CD-HIT-EST[30] was utilized to cluster and remove similar nucleotide sequences that had sequence identities of over 90%. With a high filter, we still maintain some similar genes and the CD-HIT-EST tool creates small clusters. After removal of such redundant genes, the remaining 42,266 sequences were sorted in descending order based on their CAI as depicted in S2 Appendix. Of these, 7,406 sequences with the highest CAI were selected to serve as the model dataset. Approximately 70% of the dataset was used for training (5,184 sequences), 10% for validation (741 sequences), and 20% for testing (1,481 sequences).

### Synthetic Plasmid Benchmarks

40 DNA sequences were established as a benchmark set, extracted from both studies conducted on codon optimization and gene expression evaluation of plasmids in *E. coli*. The benchmark set serves as a validation for the effectiveness of the tool on genes whose heterologous expression has been studied. The resultant coding regions of the sequences can be accessed at https://doi.org/10.5281/zenodo.5529209 and their descriptions in Table S1.

### Encoding

Encodings were created for our entire dataset using the “one-hot encoding” technique. Amino acid sequences were converted into integers and then placed into vectors that are 26 features long. At each timestep, the present amino acid is encoded into the vector as “1” while all other features are set to “0”. For example, the amino acid Alanine can be represented by a 1×26 vector in which the first element is 1 and all other elements (features) are set to 0. Features were based on Table S2. In addition, we experimented with encodings based on a Non-Linear Fisher Transform (NLFT) technique that has coded 18 features per amino acid[32].

### Model Building

Predicting an optimal synonymous codon with sequential information – the sequence of codons that surround the prediction – may yield synonymous codon selection that is more similar to the host organism. Deep learning is a technique that may be able to capture underlying patterns found in the host genome. Our model uses the BiLSTM architecture[26] which predicts synonymous codons given the input amino acid sequence. The model hyperparameters as shown in Table S3 were tuned iteratively when trained on the training and validation subsets of the dataset. L2 regularization and dropout were used to fine-tune our model and prevent overfitting. This model building overview along with a user workflow is depicted in Fig. 1.

**Fig. 1.**
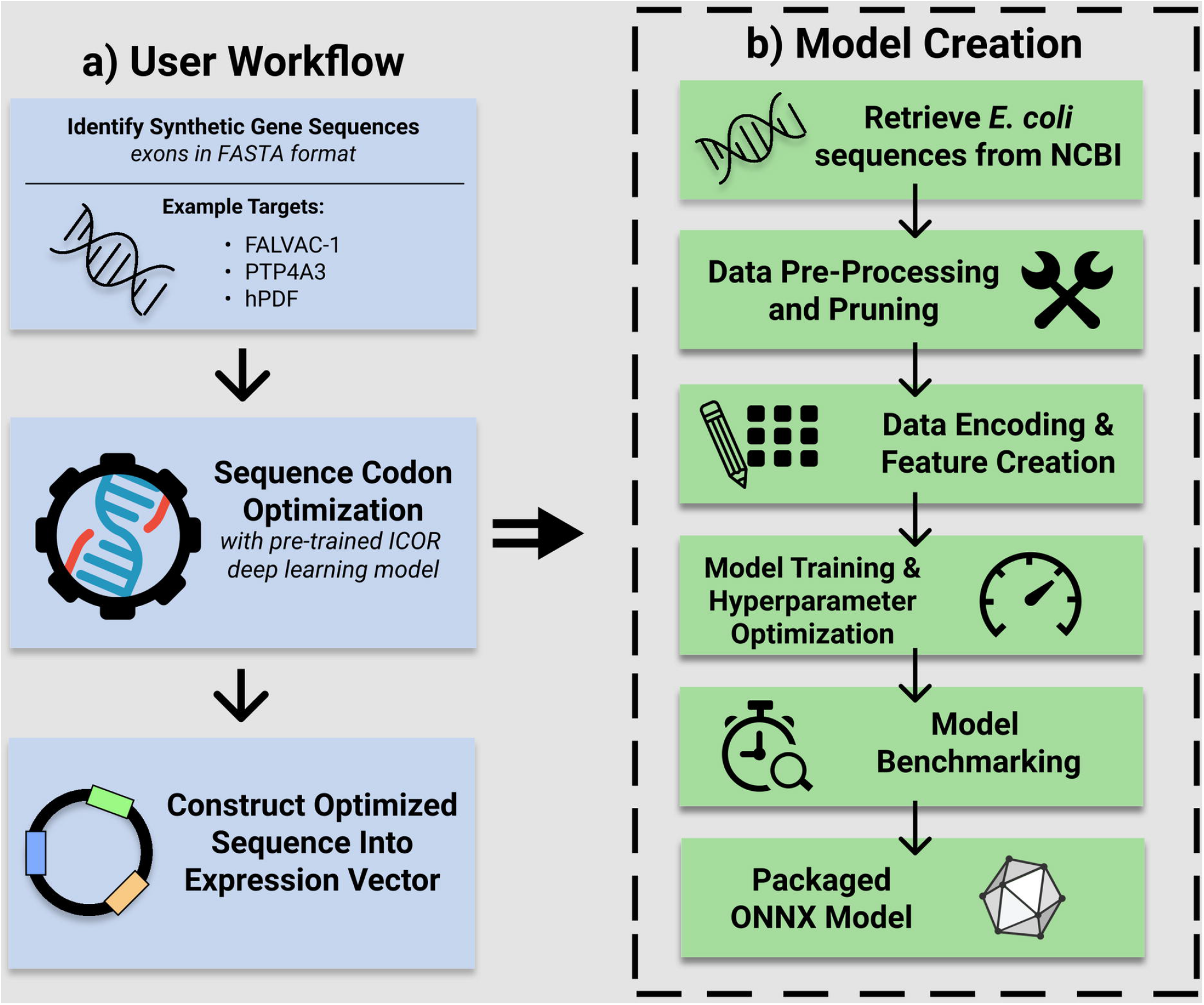
User workflow for sequence codon optimization using ICOR deep learning model with overview of model creation. (A) On the left-hand side, a user workflow towards creating a vector for heterologous expression is depicted. (B) On the right-hand side, expanding out of “sequence codon optimization” in the user workflow, the overview of the ICOR model creation is given. In a production setting, a trained and packaged model of ICOR is inferenced.

The ICOR model’s architecture consists of a 12-layer recurrent neural network as visualized in Fig. S1. Data is fed forward from the first layer—Sequence Input—to the last layer—Classification—from top to bottom. The model was trained in the MATLAB r2020b[31] on the Tesla V100 graphics card. The model was trained on 50 epochs and took 138 minutes to complete training.

### Software Architecture

The development of ICOR has two major software components for the user: ICORnet architecture and runtime scripts. The ICORnet architecture is the trained BiLSTM network. It serves as the “brain” for the codon optimization tool. By providing the amino acid sequence as an input, ICORnet can output a nucleotide codon sequence that would ideally match the codon biases of the host genome. The specifics about development are detailed in S1 Supporting Information.

With an input of sequences to be optimized, a user receives codon sequences optimized for *E. coli* expression using the ICOR runtime scripts. The runtime scripts utilize the ONNX runtime to inference the trained model.

### Statistical Analysis

The CAI is calculated using the formulae described in S1 Supporting Information. GenScript’s rare codon analysis tool[33] is utilized to calculate GC-content, CFD, negative repeat elements, and negative cis-regulatory elements. The mutational rate is quantified by conducting optimization on the test dataset, converting the optimized codons back to amino acids, and then counting the number of amino acids that varied between them. Rare codon usage was qualitatively and quantitatively compared to reference tables[34].

## Results

We use multiple previously established metrics such as the Codon Adaptation Index[35], GC-content, CFD, number of negative repeat elements, and negative cis-regulatory elements to quantify the performance of our tool. Formal definitions of these metrics are given in S1 Supporting Information.

### Codon Adaptation Index

As noted in previous studies, CAI is highly correlated with real-world expression[36]. On the test subset of 1,481 genes, we find that ICOR offered an improvement in CAI from 0.73 to 0.889 ± 0.012, or about 29.1% compared to the original sequences (Fig. 2).

**Fig. 2.**
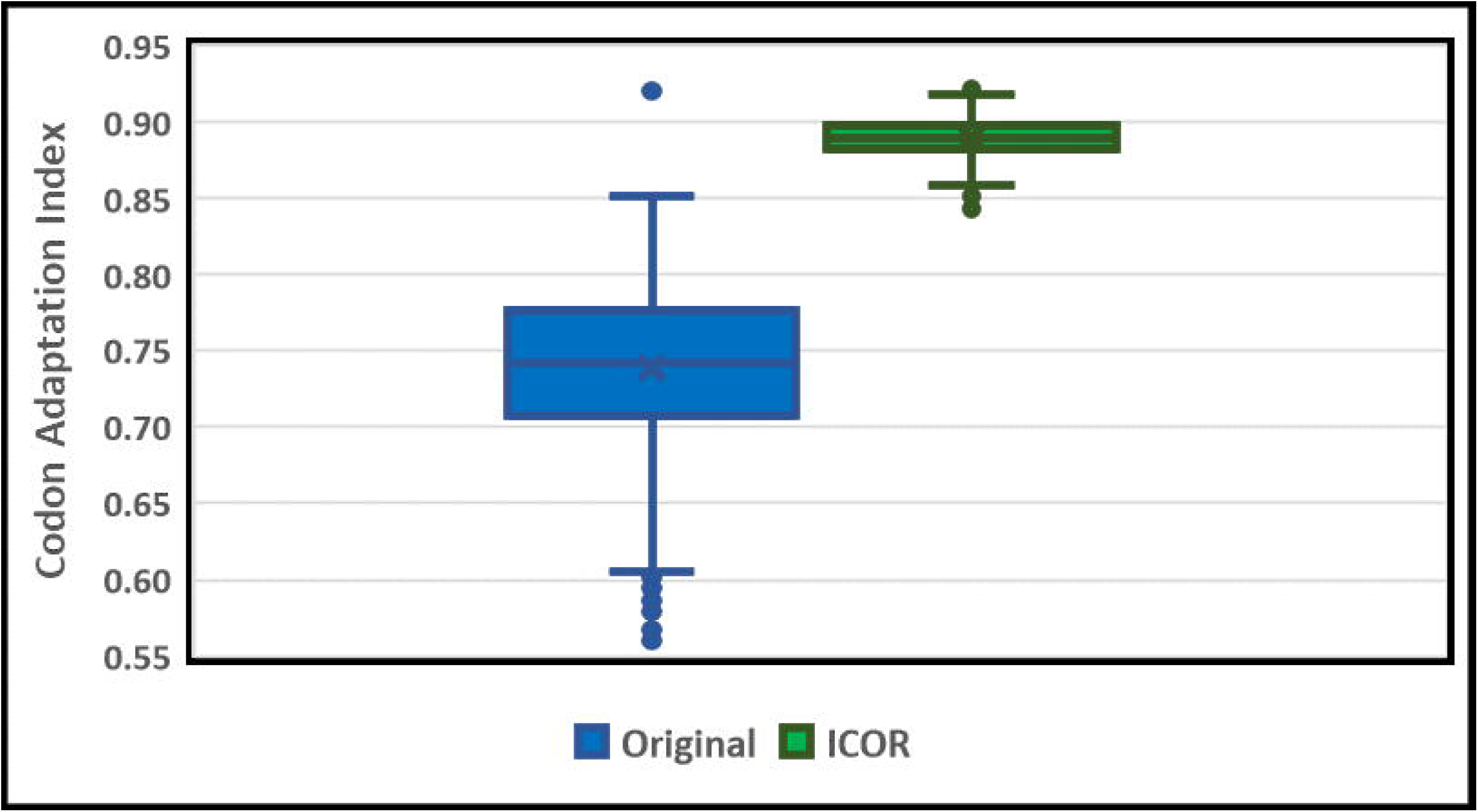
ICOR significantly improves Codon Adaptation Index (CAI) when compared to original sequences on a test subset of 1,481 genes. Box and Whisker Plot (n=1481) comparing CAI (left: original sequences, right: ICOR optimized sequences). The y-axis is the Codon Adaptation Index on a scale from 0.55 to 0.95. The open points outside of the boxes are outliers that are beyond 1.5 times the interquartile range. The horizontal divisions present in each box (from top to bottom) are the upper whisker, 3^rd^ quartile, median, 1^st^ quartile, and lower whisker.

In order to properly contextualize the performance of the developed model, six algorithms from S1 Appendix were used to optimize the benchmark set of 40 genes. These 40 genes came from a variety of origin organisms and had a mean CAI of 0.638 with a standard deviation of 0.0386. ICOR optimization yielded a mean CAI of 0.904 with a standard deviation of about 0.016, signifying a ~41.692% increase in CAI. The URC approach had a mean CAI of 0.602 and standard deviation of about 0.022. ICOR offered a ~50.21% increase in CAI compared to this approach. Finally, the BFC approach offered a mean CAI of 0.699 and a standard deviation of 0.0158. ICOR offered a ~29.32% increase in CAI compared to the BFC approach. These comparisons were statistically significant (p<0.0001) using a two-sample t-test. The mean CAI for all approaches is shown in Fig. 3.

**Fig. 3.**
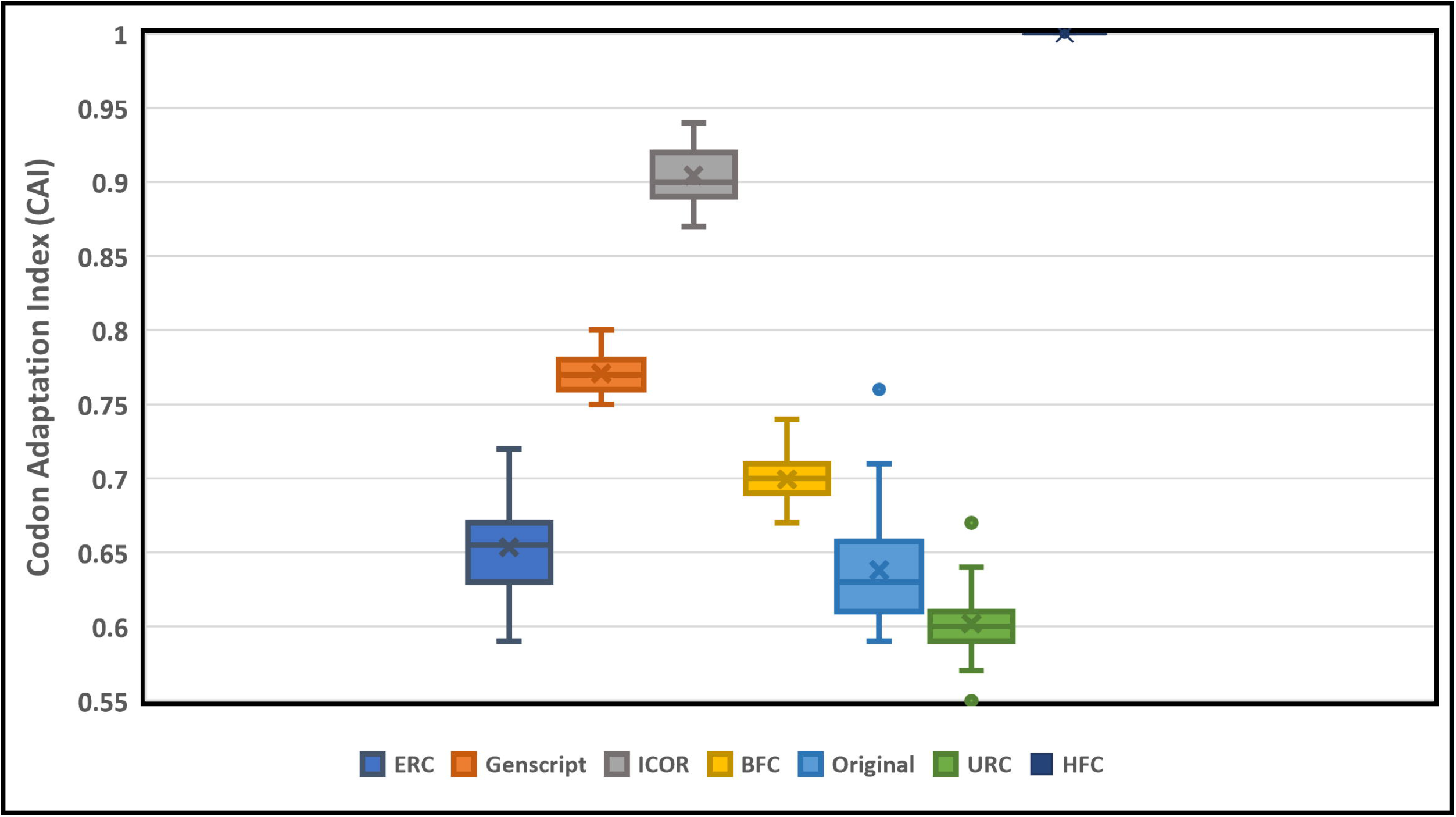
ICOR significantly improves Codon Adaptation Index (CAI) when compared to the original, BFC, URC, ERC, and GenSmart techniques. Box and Whisker Plot (n=40) comparing CAI with legend indicating the color for each optimization method.

When extrapolating such improvements to findings by dos Reis et al. on the correlation between CAI and expression for group 1 (biased) genes, real-world mRNA expression could improve by an estimated 236%[36].

### Secondary Endpoints

The secondary endpoints were quantified for each of the six codon optimization approaches on the benchmarking dataset sequences (n=40).

Ideal GC-content for recombinant genes is known to be between 30% and 70%; peaks outside of this range adversely affect transcriptional and translational efficiency[33]. ICOR, along with the other optimization techniques were all found to optimize genes within this range. The mean GC-content for each optimization technique is depicted in Fig. 4.

**Fig. 4.**
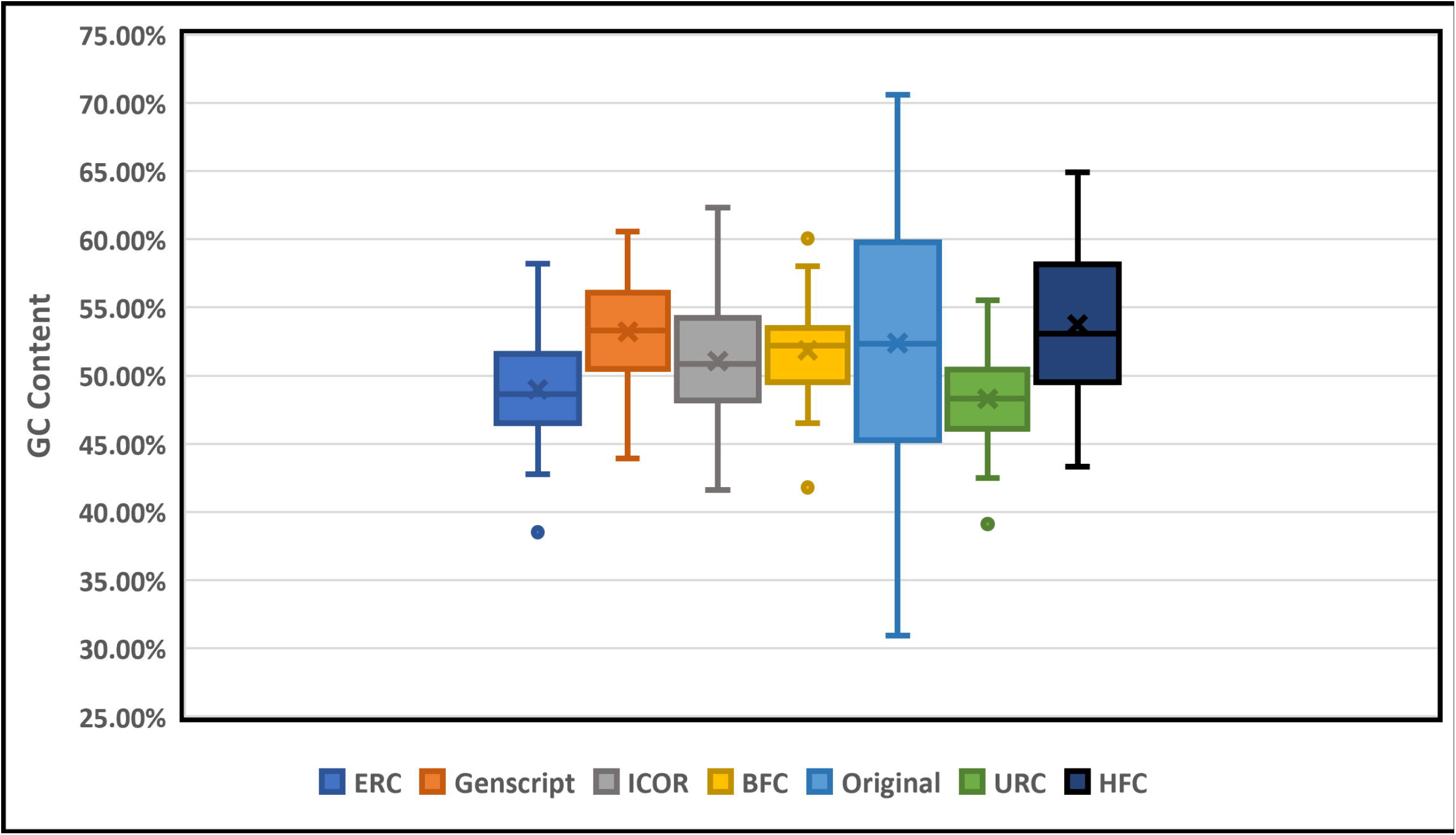
ICOR and other codon optimization techniques all maintain within optimal GC-content range. Box and whisker plot (n=40) comparing GC-content with legend indicative of each optimization method.

Genes that employ low frequency (<30% usage in the host genome) can cause a disengagement of translational machinery and reduce the efficiency of translation[33]. This rare codon frequency distribution is measured as a percentage and minimizing this value is ideal. ICOR offered a significant improvement in CFD, outperforming all the other optimization techniques tested. ICOR reduced CFD by 93.55% and 97.69% compared to the GenSmart and original sequences respectively. The improvement over GenSmart and original sequences were both statistically significant with p-values less than 0.0001. The mean CFD for each optimization technique is depicted in Fig. 5.

**Fig. 5.**
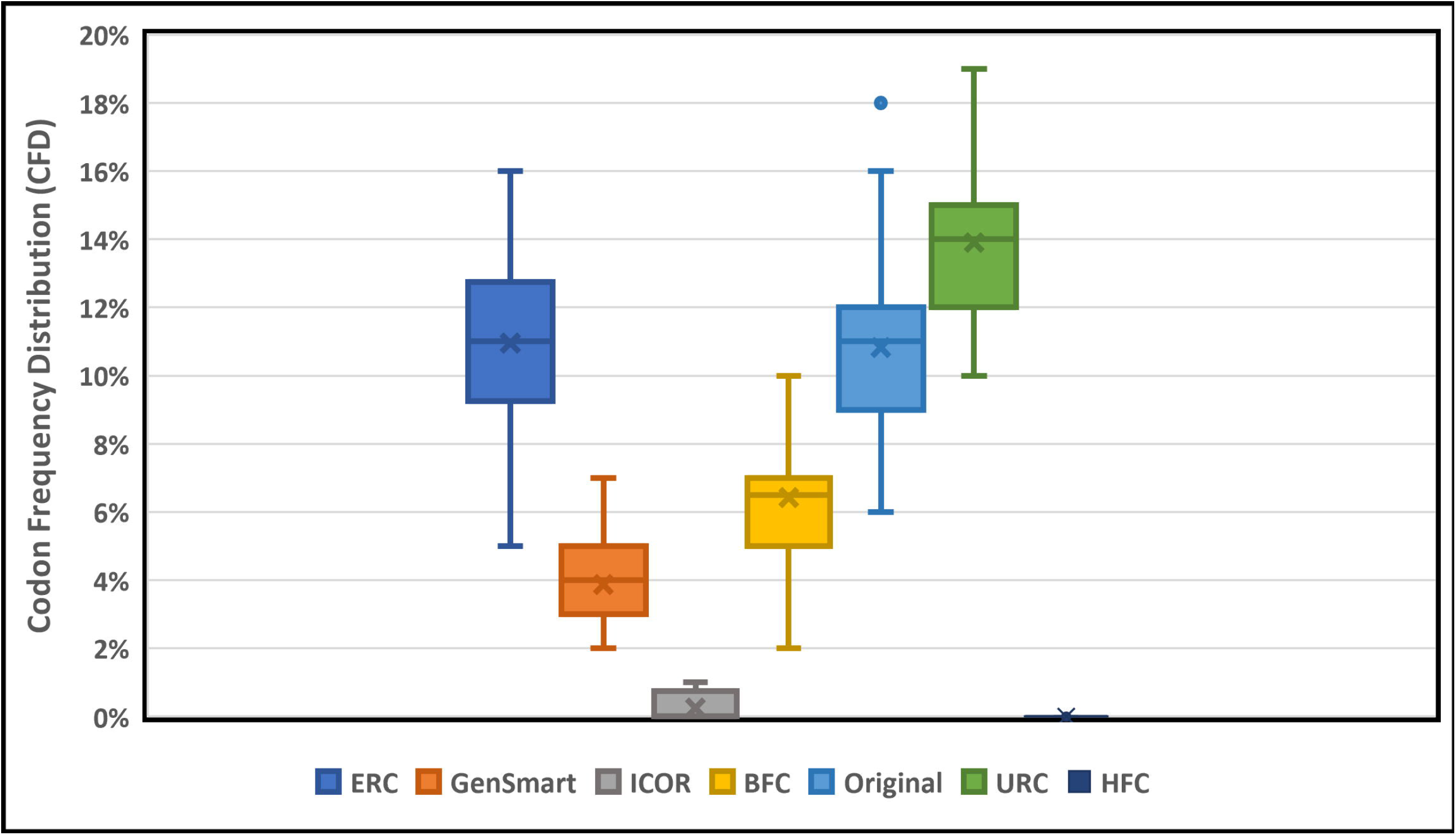
ICOR codon optimization significantly improves mean Codon Frequency Distribution (CFD). Box and whisker plot (n=40) comparing CFD with legend indicating each optimization method.

There was a difference (p=0.726) that was not found to be significant between ICOR and GenSmart in the number of negative cis-regulatory elements using a two-sided Mann-Whitney U Test for non-parametric distributions. When computing the mean change in negative cis-regulatory elements between the optimization tool and the original sequence, there is also a statistically insignificant difference between ICOR and GenSmart. This suggests that ICOR maintains equally low negative cis-regulatory elements as GenSmart while achieving a higher CAI. The mean number of negative cis-regulatory elements is depicted in Fig. 6.

**Fig. 6.**
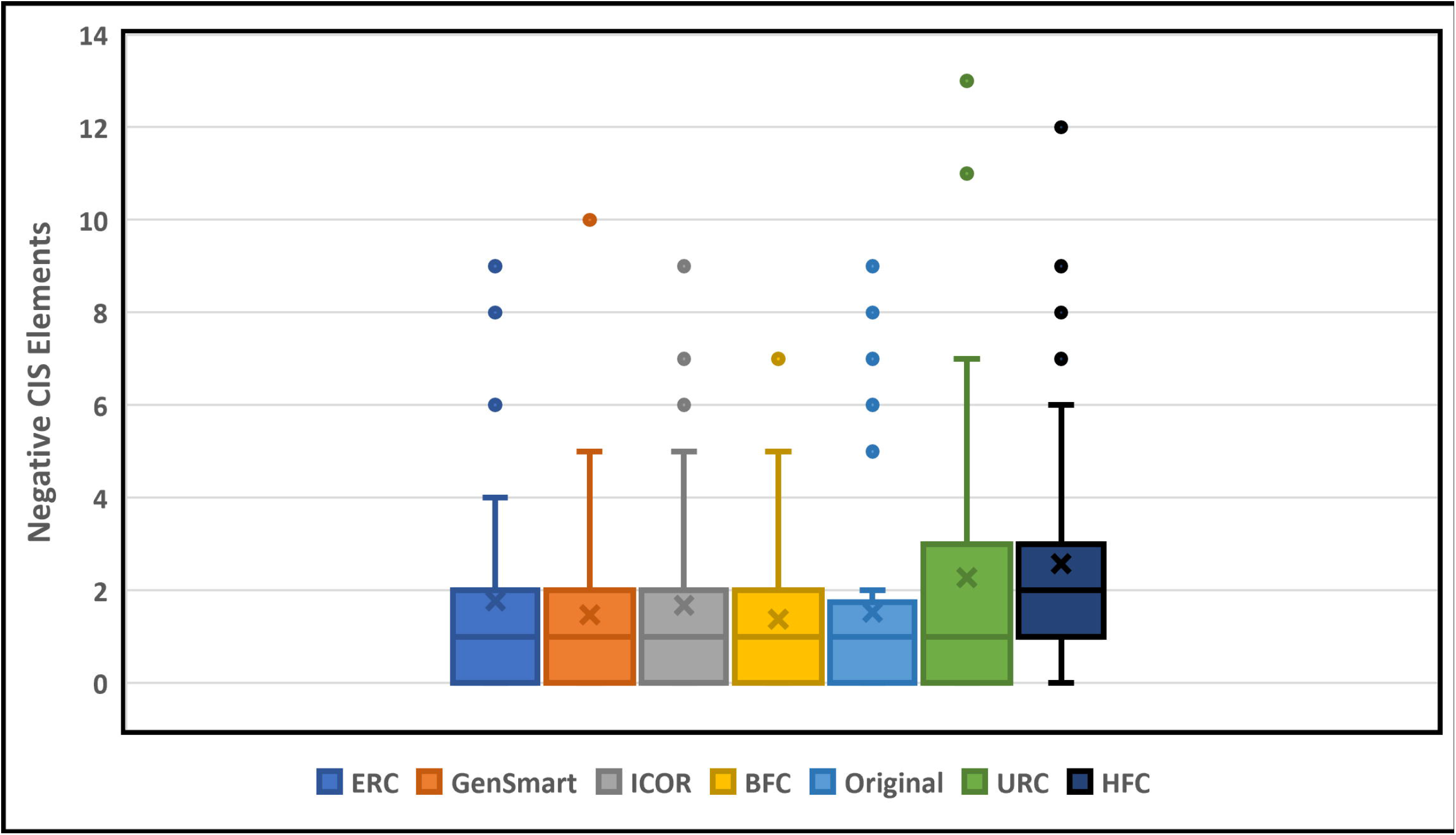
ICOR codon optimization technique yields statistically insignificant change in negative cis-regulatory elements. Box and whisker plot (n=40) comparing negative cis-regulatory elements with legend indicative of each optimization method.

There was a trending difference (p=0.1826) between ICOR and GenSmart in the number of negative repeat elements in the optimized sequences using a two-sided Mann-Whitney U Test for non-parametric distributions. This suggests that although ICOR may have higher negative repeat elements, the difference is minimal as compared to GenSmart. The mean number of negative repeat elements is depicted in Fig. 7.

**Fig. 7.**
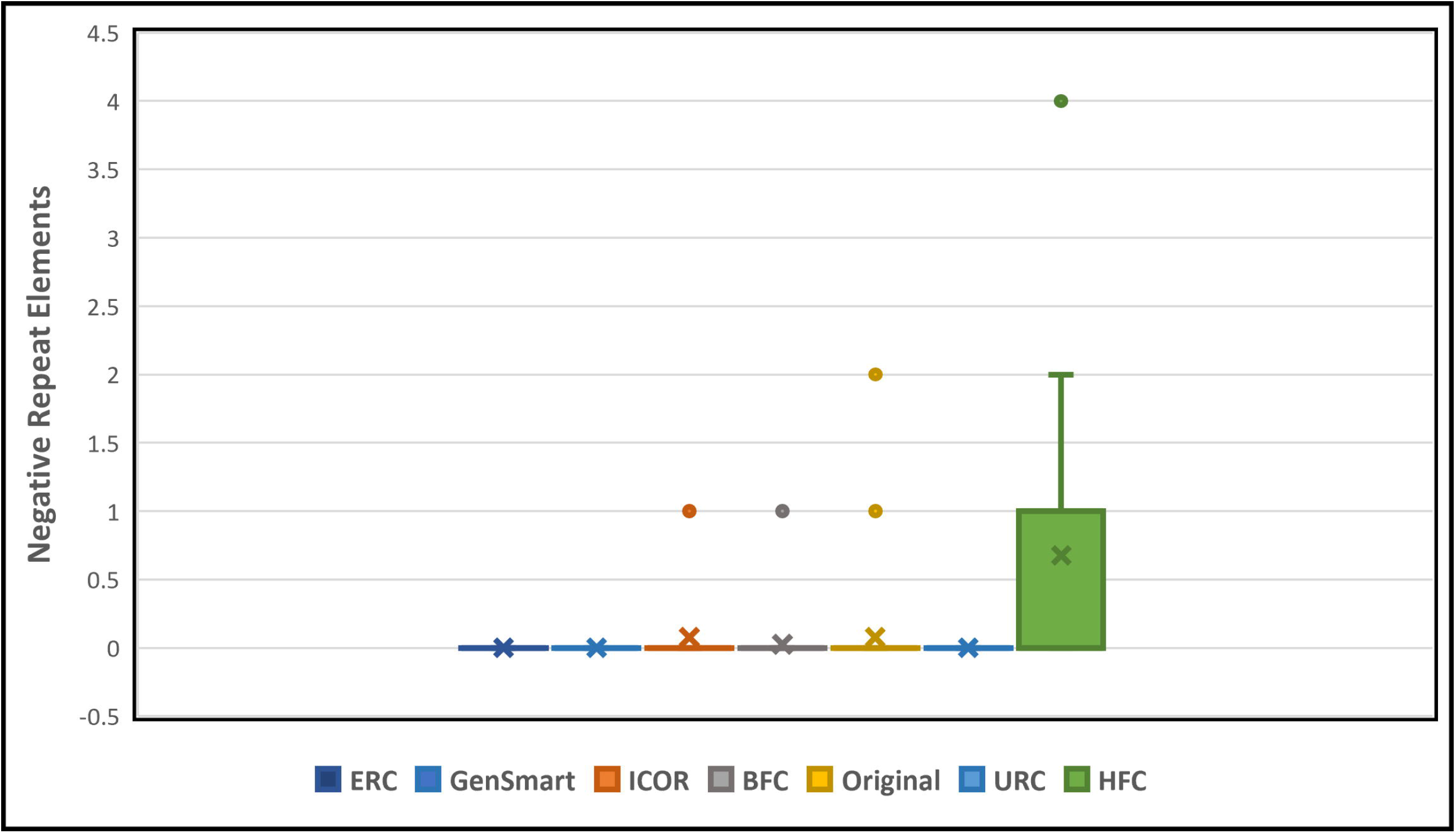
Trending difference between ICOR optimization in number of negative repeat elements. Box and whisker plot (n=40) comparing negative repeat elements with legend indicative of each optimization method.

### Optimization Run Time

The run time was calculated for the approaches where inference time could be isolated. Using a testing system as described in S1 Supporting Information, the algorithms were evaluated for run time on the benchmark set with an average length of 1687.65 nucleotides per sequence. The scores normalized to the URC approach are displayed in Table 3.

**Table 3.**
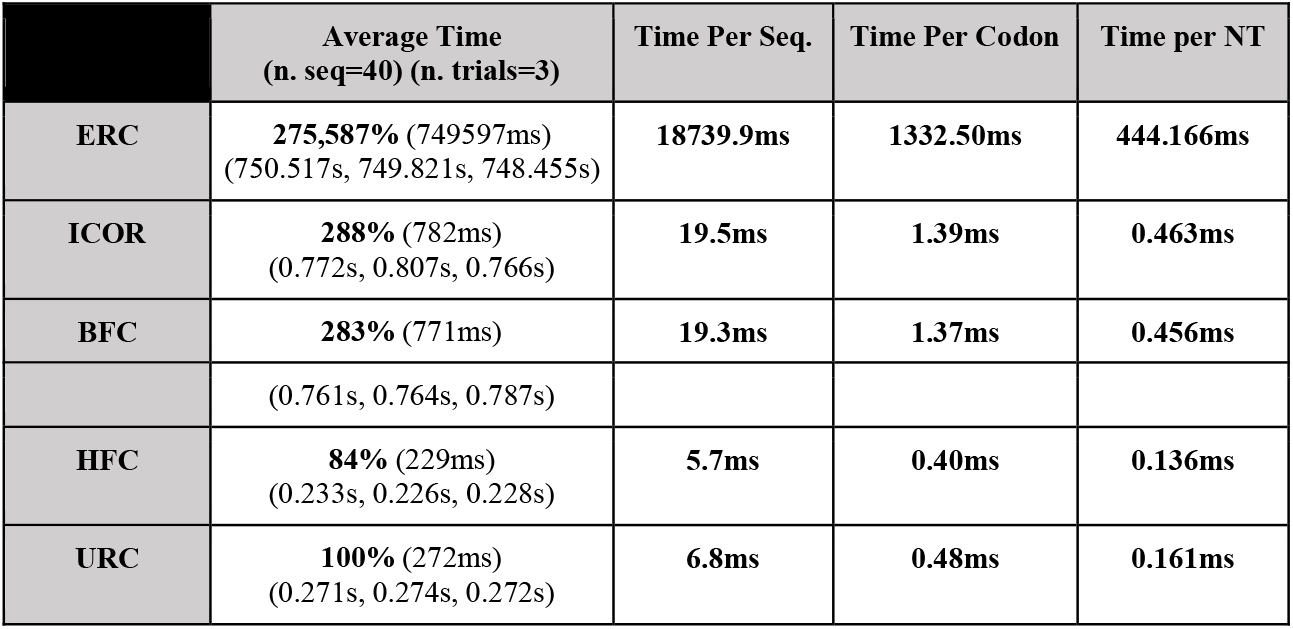
ICOR runtime comparable to the BFC, HFC, and URC approaches and significantly faster than ERC approach. Three trials were conducted with the score being an average of these three times measured to the third significant figure in milliseconds. Optimization run time is displayed with a normalized percentage to the URC approach because it represents the most naïve codon selection.

|  | Average Time<br>(n. seq=40) (n. trials=3) | Time Per Seq. | Time Per Codon | Time per NT |
| --- | --- | --- | --- | --- |
| **ERC** | **275,587%** (749597ms)<br>(750.517s, 749.821s, 748.455s) | **18739.9ms** | **1332.50ms** | **444.166ms** |
| **ICOR** | **288%** (782ms)<br>(0.772s, 0.807s, 0.766s) | **19.5ms** | **1.39ms** | **0.463ms** |
| **BFC** | **283%** (771ms) | **19.3ms** | **1.37ms** | **0.456ms** |
|  | (0.761s, 0.764s, 0.787s) |  |  |  |
| **HFC** | **84%** (229ms)<br>(0.233s, 0.226s, 0.228s) | **5.7ms** | **0.40ms** | **0.136ms** |
| **URC** | **100%** (272ms)<br>(0.271s, 0.274s, 0.272s) | **6.8ms** | **0.48ms** | **0.161ms** |

GenSmart optimization was not comparable due to its production environment with a queue of jobs.

### Gene Mutations

The deep learning model in this study does not have strict rules in place regarding codon usage: it is not explicitly given a “codon to amino acid dictionary.” At least one point mutation would arise if a codon prediction is replaced with a non-synonymous choice (e.g., CAA to GAA). On the test dataset of 1,481 genes, our model yielded a 0.00% mutational rate and did not predict non-synonymous codons. Thus, it was found that gene mutations are not present in the ICOR codon optimization technique.

During our testing, it was found that encoding techniques made a significant difference in learning these initial codons to amino acid pairings. The One-Hot Encoding technique offered approximately a 10% improvement in matching prediction of the host codon over NLFT during model training.

## Conclusions

In this paper, we introduce ICOR, a codon optimization tool that uses recurrent neural networks towards improving heterologous expression for synthetic genes. We find that deep learning is a particularly effective codon optimization method because it learns codon usage bias in tandem with a codon’s surrounding context, thus allowing it to make predictions using the patterns and subsequences in which synonymous codons are used in genes. While previous research ranges from selecting high-frequency codons, to eliminating secondary structures, to machine learning models and convolutional neural network architectures, we use the RNN architecture which has the ability to take sequential information into account. The RNN can thereby use underlying patterns in genes to inform codon selection that may be more similar to that of the host genome.

Using this approach, we built the ICOR model using a large dataset of 7,406 non-redundant genes from the *E. coli* genome. Having a non-redundant dataset was of vital importance in this study as many *E. coli* genomes across various strains contain similar genes. We used the CD-HIT-EST server to overcome this issue because a model trained on redundant data would yield codon selection that is biased to common *E. coli* genes only. Further, this helped reduce the necessary compute resources required for deep learning. The ICOR model encodes gene sequences using Natural Language Processing techniques, demonstrating their effectiveness in this task.

The statistical data analysis uses the CAI, GC-content, CFD, negative repeat elements, and negative cis-regulatory elements as metrics to compare ICOR to other solutions. Although ICOR demonstrates improvements in these metrics, signifying increased protein expression, they are not the only comprehensive statistics to predict gene expression. Recent research has shown that creating models for measuring translation dynamics is possible[37], and such research can be applied to the results of our study. Modeling elongation rate, tRNA adaptation index, and other metrics may provide further valuable insight into the results of our tool.

In this research, six codon optimization techniques were evaluated. Of these, the ERC algorithm attempts to illustrate the efficacy of an exhaustive optimization in which all potential sequences are generated. Due to computational limitations, a truly exhaustive search method in which every possible gene sequence is evaluated cannot be efficacious. Given an average sequence length of 562.55 amino acid and an average of 2.91 potential codons, over 1.01 × 10^8^ possible sequences would exist, requiring an infeasible amount of time (over 524,000 hours) to calculate. Such a method would operate under the assumption that there is a single metric that requires optimization. However, our review suggests that optimization may instead require a genome-wide understanding of codon usage to yield optimal expression. The HFC algorithm illustrates the implications of a “CAI = 1.0” optimization strategy which the results point towards increased negative cis-regulatory elements and negative repeat elements. In addition, our review suggests that this method may result in plasmid toxicity.

ICOR codon optimization is competitive and can be applied directly in synthetic gene design. Synonymous codons can be optimized to increase resultant protein expression. Thus, the efficiency of production improves, potentially decreasing the cost of *E. coli* recombinantly-produced products. Currently, the ICOR tool can be accessed through an open-source software package at https://doi.org/10.5281/zenodo.5529209, however, we would like to build an API to improve accessibility in the future.

Although our model is based on *E. coli* genomes, it may be possible to apply our methodology to other organisms such as yeast and mammalian cells in future research. A transfer learning approach may allow us to preserve our pre-trained model and adapt it to other host cells. Additionally, we would like to add the ability for our model to optimize other regions of a gene such as promoter sequences. Research aimed at analyzing what sub-sequence properties are learned by the model to make predictions may be biologically relevant.

Finally, experimental results for our method are not included. This is relevant. As a contribution to bioinformatics and machine learning, biological results would only be useful to demonstrate our ability to synthesize specific DNA sequences, which is outside the scope of this paper. The efficacy of these sequences would only feedback into our machine learning workflow and not fundamentally change the process as outlined.

## Availability and requirements

Project name: ICOR: Improving Codon Optimization with Recurrent neural networks

Project home page: https://github.com/Lattice-Automation/icor-codon-optimization

Operating system(s): Platform independent

Programming language: Python

Other requirements: Python 3.9.4 or newer

License: MIT

Any restrictions to use by non-academics: license needed

## Declarations

### Abbreviations

*E. coli*: *Escherichia coli*
RNNs: Recurrent neural networks
LSTM: Long short-term memory
BiLSTM: Bidirectional LSTM
CAI: Codon adaptation index
CFD: Codon frequency distribution
NCBI: National Center for Biotechnology Information
NLFT: Nonlinear Fisher Transformation
ERC: Extended Random Choice
BFC: Background Frequency Choice
URC: Uniform Random Choice

## Ethics approval and consent to participate

Not Applicable

## Consent for publication

Not Applicable

## Availability of data and material

The training dataset was derived from the National Center for Biotechnology Information’s (NCBI) GenBank database [https://www.ncbi.nlm.nih.gov/assembly/?term=escherichia+coli]

The benchmark sequence datasets used to compare codon optimization approaches are available in the ICOR repository [doi:10.5281/zenodo.5529209] and see Reference [29]. Please see S1_Table for public references used to derive the training dataset.

## Authors’ contributions

RJ, AJ, and DD were involved in determining the datasets and methods of comparison for codon optimization tools. RJ, AJ, DD, KL were involved in the writing of the manuscript and in the interpretation of the results. RJ, KL, and EM were involved in administration. All authors edited, revised, read and approved the final manuscript.

## Acknowledgements

Not Applicable

## Additional Files

1. File name: S1_Fig.pdf
  a. Title: ICOR deep learning model architecture
  b. Description: An overview of the 12-layer recurrent neural network model trained in this study.
2. File name: S1_Appendix.docx
  a. Title: Codon optimization methods overview
  b. Description: An overview of the four codon optimization methods built and defined in this study.
3. File name: S1_SupportingInformation.docx
  a. Title: Supporting details for testing methods and computational metrics
  b. Description: Specifications for the system used in the benchmarking of the software tool along with supporting formulae used in the validation of the software tool performance.
4. File name: S1_Table.docx
  a. Title: Description of 40 genes used to benchmark codon optimization methods
  b. Description: Average amino acids/codons in sequence: 562.55. Average length/nucleotides of sequence: 1687.65. Genes are sorted by the number of base pairs from smallest to greatest.
5. File name: S2_Appendix.docx
  a. Title: Training dataset specifications
  b. Description: A histogram displaying the distribution of the codon adaptation index of genes utilized in the ICOR training dataset.
6. File name: S2_Table.docx
  a. Title: Amino acid integer lookup table
  b. Description: A list of the encoding used for converting amino acids to numerical, integer values.
7. File name: S3_Table.docx
  a. Title: Hyperparameter variables are used to finetune the results of the model
  b. Description: (A) In the left-hand column, 8 hyperparameters are listed. (B) In the righthand column, their respective settings are given based on the testing and fine-tuning conducted in this research. Our iterative tuning methods reach the following hyperparameters, which prevent the possibility of over/underfitting while maintaining high performance.

