## Supplementary material for "ICOR: Improving codon optimization with recurrent neural networks": S1 Table

| Gene Name | Gene Description & Selection Reasoning |
|---|---|
| BIRC5<br>**aa: 166**<br>**bp: 498** | BIRC5 (Baculoviral IAP repeat-containing 5) is a multitasking protein that has roles in preventing apoptosis and cell proliferation (PubMed:9859993, PubMed:21364656, PubMed:20627126, PubMed:25778398, PubMed:28218735). In Reference Paper, BIRC5 expression levels were analyzed in E. coli. Sequence optimization (generated by using a sliding window → narrowed down) was used for this human gene to be expressed in E. coli. |
| CAV1<br>**aa: 179**<br>**bp: 537** | CAV1 (Caveolin 1, caveolae protein) is involved in promoting cell cycle progression. It's a gene of interest because it is a tumor suppressor candidate making it potentially useful in recombinant expression. In Reference Paper, CAV1 expression levels were analyzed in E. coli. |
| PTP4A3<br>**aa: 187**<br>**bp: 561** | Protein tyrosine phosphatase 4A3 (PTP4A3/PRL-3) is a well-documented protein that can be produced recombinantly in E. coli. This protein has been identified as a potential target to treat some cancers. It has been studied in (codon optimization papers). |
| PA<br>**aa: 187**<br>**bp: 561** | Polymerase acidic protein (PA) plays a role in viral RNA transcription and replication. It is from the Influenza A virus. It has been studied in (codon optimization papers). |
| hPDF<br>**aa: 205**<br>**bp: 615** | Human peptide deformylase (hPDF) is a target for cancer therapeutics. However, its expression is not very efficient in E. coli. This serves as a good target for benchmarks as past studies have noted the valuable potential of codon optimization of this gene. |
| EMG1<br>**aa: 245**<br>**bp: 735** | EMG1 (Protein: nucleolar protein homolog (S. cerevisiae)) is a human-based recombinant protein expressed in *E. coli*. This gene encodes a protein that methylates pseudouridine and is an essential eukaryotic protein. In Reference Paper, EMG1 expression levels were analyzed in *E. coli*. |
| CDK1<br>**aa: 298**<br>**bp: 894** | CDK1 (Cyclin Dependent Kinase 1) codes for a protein which is essential for G1/S and G2/M phase transitions. It has been used in calculating prognostic value in human cancer (Reference Paper). Higher expression levels correlated with a more-advanced tumor. Another reference paper analyzing the importance of this gene looked at expression and structure through recombinant expression in *E. coli* (Reference Study). |
| CD80<br>**aa: 307**<br>**bp: 921** | Cd80 (Mus musculus CD80 antigen) is a protein-coding gene whose protein, once activated, induces T-cell proliferation and cytokine production. In Reference Paper, Cd80 expression levels were analyzed in E. coli. |

| | |
|---|---|
| TAS2R10<br>**aa: 308**<br>**bp: 924** | Taste receptor, type 2, member 10 (TAS2R10) encodes proteins expressed in taste receptor cells. In Reference Paper, TAS2R10 expression levels were analyzed in *E. coli*. |
| PIM1<br>**aa: 314**<br>**bp: 942** | Pim-1 oncogene (PIM1) encodes a protein that plays a role in signal transduction in blood cells. The gene is expressed primarily in B-lymphoid and myeloid cell lines. It is found to be overexpressed in hematopoietic malignancies and in prostate cancer. In Reference Paper, PIM1 expression levels were analyzed in *E. coli*. |
| FALVAC-1<br>**aa: 324**<br>**bp: 972** | FALVAC-1 is a vaccine against Plasmodium Falciparum. This is a good benchmark as a synthetic gene of codon optimization tools because it is a real candidate vaccine that is recombinantly produced (Reference Paper). |
| CREB1<br>**aa: 328**<br>**bp: 984** | CREB1 (cAMP responsive element binding protein 1) mutations can increase risk for certain diseases. Its most common variant is actually from a tumor. It is from a family of transcription factors that is expressed in the brain. In Reference Paper, CREB1 expression levels were analyzed in *E. coli*. |
| JUN<br>**aa: 332**<br>**bp: 996** | JUN (Jun Proto-Oncogene, AP-1 Transcription Factor Subunit) can often be mutated to cause a cell to become a tumor cell. JUN is the "putative transforming gene" of avian sarcoma virus 17 (Reference). It was expressed as a "transcription factor" in this Reference Paper in *E. coli*; its expression was measured in vivo. |
| CSNK1A1<br>**aa: 338**<br>**bp: 1014** | CSNK1A1 (Casein Kinase 1 Alpha 1) belongs to the protein kinase superfamily and is involved in phosphorylating numerous proteins involved in cellular functions. Recombinant human CSNK1a1 protein, fused to His-tag at N-terminus,was expressed in *E.coli* by CSNK1A1, 1-337aa Human, His tag, E.coli (GWB-ATG0D4). |
| MAPK1<br>**aa: 361**<br>**bp: 1083** | Mitogen-Activated Protein Kinase 1 (MAPK1) is a part of the MAP kinase signal transduction pathway. It is of special interest because in this Reference Paper, when expressed in *E. coli*, there was a significant difference in the protein yield for their wild-type and optimized genes: 24.3 and 11.5 mg/L respectively. |
| OPRM1<br>**aa: 401**<br>**bp: 1203** | "This gene encodes one of at least three opioid receptors in humans; the mu opioid receptor (MOR). The MOR is the principal target of endogenous opioid peptides and opioid analgesic agents such as beta-endorphin and enkephalins. The MOR also has an important role in dependence to other drugs of abuse, such as nicotine, cocaine, and alcohol via its modulation of the dopamine system" (Study). In Reference Paper, OPRM1 expression levels were analyzed in *E. coli*. |
| LAMP1<br>**aa: 418**<br>**bp: 1254** | Lysosomal-associated membrane protein 1 (LAMP1) is a protein coding gene that is associated with Chediak-Higashi Syndrome and Gaucher's Disease.This is a gene of interest because it may play a role in tumor cell metastasis. It was |

| | |
|---|---|
| | expressed as a "transcription factor" in this Reference Paper in *E. coli*; its expression was measured in vivo. |
| NGFR<br>**aa: 428**<br>**bp: 1284** | Nerve growth factor receptor (NGFR) is a protein coding gene associated with important pathways within the cell. Diseases associated with NGFR include Prurigo Nodularis and Infiltrative Basal Cell Carcinoma. In Reference Paper, NGFR expression levels were analyzed in *E. coli*. |
| GSK3B<br>**aa: 434**<br>**bp: 1302** | GSK3B (Glycogen Synthase Kinase 3 Beta) encodes a protein that is a negative regulator of glucose homeostasis. Mutations that lead to defects in this gene are associated with Parkinson disease and Alzheimer disease. This is a gene of interest because it is found to be an important enzyme in glycogen metabolism. In Reference Paper, GSK3B expression levels were analyzed in *E. coli*. |
| CLN3<br>**aa: 439**<br>**bp: 1317** | CLN3 (Ceroid-lipofuscinosis, neuronal 3) is a gene that encodes a protein involved in lysosomal function. This is an interesting gene of study because it can cause neurodegenerative diseases once mutated (Batten disease). In Reference Paper, CLN3 expression levels were analyzed in *E. coli* which makes it a good benchmark gene to compare to. |
| MAPKAPK5<br>**aa: 472**<br>**bp: 1416** | Mitogen-activated protein kinase-activated protein kinase 5 (MAPKAPK5) is a gene that encodes for a tumor suppressor. It is activated by the MAPK1 kinase and also plays an important role in the MAP kinase signal transduction pathway. It is of special interest because in this Reference Paper, when expressed in *E. coli*, there was a significant difference in the protein yield for their wild-type and optimized genes: 14.9 and 80 mg/L respectively. |
| SMARCD1<br>**aa: 475**<br>**bp: 1425** | SWI/SNF related, matrix associated, actin dependent regulator of chromatin, subfamily d, member 1 (SMARCD1) is a gene that encodes a protein which is thought to regulate transcription. It has been found to inhibit certain human glioblastoma cells, sparking interest in the overexpression of this gene. In Reference Paper, SMARCD1 expression levels were analyzed in *E. coli*. |
| AKT1<br>**aa: 481**<br>**bp: 1443** | AKT serine/threonine kinase 1, naturally found in humans, encodes one of three kinases which are referred to as protein kinase B alpha, beta, and gamma. In the Reference Paper, AKT1 was found to have a reduction in optimized heterologous expression by 10% compared to the wild-type gene. This makes it a particularly interesting benchmark gene to gauge the performance of codon optimization tools. |
| LCK<br>**aa: 510**<br>**bp: 1530** | Lymphocyte-specific protein tyrosine kinase (LCK) is a gene that encodes a protein that is a key signaling molecule "...in the selection and maturation of developing T-cells" (Reference). In Reference Paper, LCK expression levels were analyzed in *E. coli*. It was found that LCK showed a 500-fold increase in mRNA transcripts for the sequence-optimized gene. |

| | |
|---|---|
| RPS6KB1<br>**aa: 526**<br>**bp: 1578** | Recombinant Human Ribosomal protein S6 kinase beta-1 (RPS6KB1) promotes cell proliferation and protein synthesis. It has been extensively studied in yeast and *E. coli* expression systems. |
| PAK1<br>**aa: 546**<br>**bp: 1638** | p21 protein (Cdc42/Rac)-activated kinase 1 (PAK1) is a gene that encodes a family member of p21-activating kinases. Overexpression of this gene can result in cancer growth and mutations in this gene are associated with diseases such as macrocephaly. In Reference Paper, PAK1 expression levels were analyzed in *E. coli*. |
| PLK1<br>**aa: 604**<br>**bp: 1812** | Polo like kinase 1 (PLK1) is a human gene that has been targeted in cancer therapy. A reduction in PLK1 levels is associated with the inhibition of cell proliferation and induced apoptosis. In Reference Paper, PLK1 expression levels were analyzed in *E. coli*. |
| PEA<br>**aa: 613**<br>**bp: 1839** | Pseudomonas aeruginosa exotoxin A (PEA) is an important pathogenic factor 1. It retains high immunogenicity even after detoxification, enabling its use as vaccine adjuvants and vaccine carriers. This is a good benchmark as it can be produced for a low-cost at a large-scale in E. coli. Further, the (Reference Paper) finds that codon optimization enhances expression of PEA in E. coli -- thus, if the tool presented in this research can achieve similar/better results than the paper and/or other approaches, it will be considered an improvement. |
| TAP1<br>**aa: 749**<br>**bp: 2247** | Transporter 1, ATP-binding cassette, sub-family B (MDR/TAP) (TAP1) encodes a protein known to be involved in molecular transport and drug resistance. In Reference Paper, TAP1 expression levels were analyzed in *E. coli*. |
| NOC2L<br>**aa: 750**<br>**bp: 2250** | Nucleolar complex associated 2 homolog (S. cerevisiae) (NOC2L) can control major aspects of transcriptional regulation. It is an RNA/Ribosomal protein. In Reference Paper, NOC2L expression levels were analyzed in *E. coli*. |
| UBTF<br>**aa: 765**<br>**bp: 2295** | Upstream Binding Transcription Factor (UBTF) encodes a protein important for ribosomal RNA transcription. The UBTF studied originates from human genes. UBTF is well-known as a recombinant protein and has been noted as useful as a blocking peptide for certain antibodies. |
| BRAF1<br>**aa: 767**<br>**bp: 2301** | BRAF1 (v-raf murine sarcoma viral oncogene homolog B1) is a protein kinase that transduces mitogenic signals from the cell membrane to the nucleus. This kinase phosphorylates MAP2K1 which activates the MAP kinase signal pathway (PubMed:21441910, PubMed:29433126). This gene is a gene of interest because it is frequently mutated and allows a cell to become a tumor cell. In Reference Paper, BRAF1 expression levels were analyzed in E. coli. |
| FGFR4<br>**aa: 803** | FGFR4 (Fibroblast growth factor receptor 4) comes under the kinases category and the protein encoded by this gene is a tyrosine kinase and cell surface receptor |

| | |
|---|---|
| **bp: 2409** | for fibroblast growth factors. Diseases associated with FGFR4 include Prostate Cancer and Rhabdomyosarcoma. In Reference Paper, FGFR4 expression levels were analyzed in *E. coli*. |
| LEMD3<br>**aa: 912**<br>**bp: 2736** | LEM domain containing 3 (LEMD3) is a human gene that is of interest. Mutations in this gene have been associated with osteopoikilosis, Buschke-Ollendorff syndrome and melorheostosis. It was expressed as a "membrane protein" in this Reference Paper in E. coli; its expression was measured in vivo. |
| MMLP3<br>**aa: 945**<br>**bp: 2835** | Proteins associated with the MMP family are known to be involved in the breakdown of extracellular matrices in physiological processes within the cell. The MMP3 or MMLP3 gene encodes an enzyme that degrades glycoproteins such as fibronectin. It has been studied in (codon optimization papers). |
| CEBPZ<br>**aa: 1055**<br>**bp: 3165** | CEBPZ (CCAAT/enhancer binding protein zeta) is a human protein-encoding gene that plays a role in responding to environmental stimuli (heat). In Reference Paper, CEBPZ expression levels were analyzed in *E. coli* which makes it a good benchmark gene to compare to. |
| KIF11<br>**aa: 1057**<br>**bp: 3171** | KIF11 (Kinesin family member 11) was expressed recombinantly in *E. coli* in this study. "This gene encodes a motor protein that belongs to the kinesin-like protein family. Members of this protein family are known to be involved in various kinds of spindle dynamics." (KIF11 Gene (Protein Coding)). |
| NPR1<br>**aa: 1062**<br>**bp: 3186** | NPR1 (Natriuretic Peptide Receptor 1) is a gene that encodes a peptide receptor that is located in the kidney, lungs, and adipocytes. NPR1 is associated with diseases including congestive heart failure and malt worker's lung. In Reference Paper, NPR1 expression levels were analyzed in *E. coli*. |
| FLT1<br>**aa: 1339**<br>**bp: 4017** | FLT1 (Fms-related tyrosine kinase 1) is a gene that encodes a member of the vascular endothelial growth factor receptor family. In Reference Paper, FLT1 expression levels were analyzed in *E. coli*. |
| PDCD11<br>**aa: 1872**<br>**bp: 5616** | Programmed Cell Death 11 (PDCD11) is a useful binding protein that colocalizes with U3 RNA (MIM 180710) in the nucleolus and is required for rRNA maturation and generation. It is important because, as a plasma protein, it is within the nucleolus of the cell and is a ribosomal protein. In Reference Paper, PDCD11 expression levels were analyzed in *E. coli*. |
