## Supplementary figures and images for "ICOR: Improving codon optimization with recurrent neural networks"

### S1 Figure

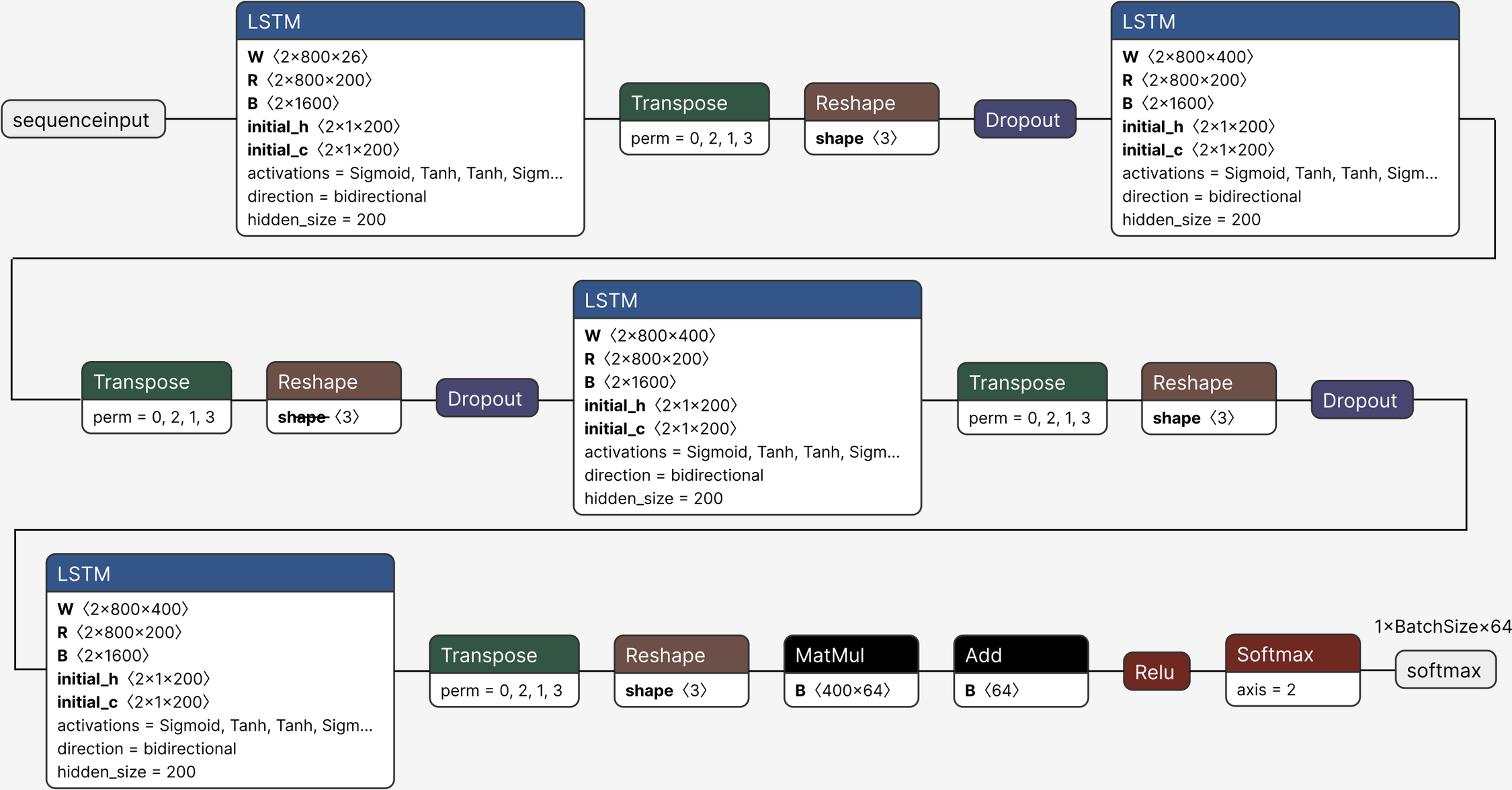
