## Supplementary material for "ICOR: Improving codon optimization with recurrent neural networks": S1 Appendix

**Uniform Random Choice Approach**

The uniform random choice (URC) utilizes a look-up table that defines amino acids and their corresponding codons. Sample:

*"A ": ["GCT GCC GCA GCG"],*

*"R": ["CGT CGC CGA CGG AGA AGG"],*

*"N": ["AAT AAC"],*

For each amino acid present in the benchmark sequence, the URC algorithm will randomly select a corresponding codon.

**Background Frequency Choice Approach**

The Background Frequency Choice (BFC) approach utilizes a look-up table that defines amino acids and their corresponding codons, along with probabilities for how frequently these codons appear. Sample:

*"A": (["GCG", "GCA", "GCT", "GCC"], [0.34, 0.22, 0.17, 0.27]),*

*"N": (["AAT", "AAC"], [0.46, 0.54]),*

For each amino acid present in the benchmark sequence, the naive approach uses the *np.random.choice* method to select a corresponding codon given the probabilities that they occur in E. coli. Such probabilities were normalized to sum to exactly 1.0.

**ICOR Approach**

The ICOR approach utilizes an RNN that contains BiLSTM layers. The model is trained on high-frequency E. coil genomes themselves. As a result, the model is able to learn the codon usage distributions of high-frequency sequences, while still keeping knowledge of subsequences and patterns for how the codons are used. For each of the 40 sequences, inference was performed using the ICOR model in MATLAB. The output sequences were saved in the FASTA format.

**Extended Random Choice Approach**

The extended random choice (ERC) approach generates 10,000 sequences by iterating through the original sequence and converting each amino acid to a random corresponding codon that codes for that amino acid (uniform random choice). With this new output, the CAI is calculated. The ERC approach returns the sequence generated that has the highest CAI. The weights to calculate CAI are defined along with the full algorithm code in the ICOR GitHub repository which can be accessed at https://doi.org/10.5281/zenodo.5529209.

**Highest Frequency Choice Approach**

The highest frequency choice (HFC) approach utilizes a look-up table that lists each amino acid and its corresponding codon that yields a CAI equal to 1.0. For each amino acid present in the benchmark sequence, the HFC approach replaces it with the highest-frequency codon. The resulting sequence has a CAI of 1.0.
