## Supplementary material for "ICOR: Improving codon optimization with recurrent neural networks": S1 Supporting Information

**Key Steps of Development:**

1. "Fastalator" loads DNA sequences from FASTA files and converts them into structures containing amino acid sequences and the original DNA sequence. It can account for DNA sequences that do not have 100% confidence (occurs during genome sequencing) by using IUPAC probability estimators.
2. "CreateTrainingData" and "CreateTrainingDataNLF" are two scripts that encode the amino acid data as vectors for the ICORnet RNN model. One uses One-Hot Encoding while the other uses Non-Linear Fisher Transform (NLFT).
3. "trainNet" trains the ICORnet model on the data imported. It provides options to adjust hyperparameter optimization and network architecture. It outputs a network object which is used for prediction of synonymous codons.

**System specifications and environment setup:**

The ICOR model was built and trained on a testing system with specifications below. The benchmarking for optimization run-time was conducted on this machine as well.

- Intel® Core™ i7-10700 Processor @ 2.90 GHz
- 32 gigabytes, 2933-MHz RAM
- 3,500/3,200 MB/s read/write SSD
- NVIDIA Turing™ GeForce RTX™ 2060 (6GB VRAM)

The software specifications on which the ICOR tool was built are specified below, along with the environment setup for the benchmarking tests.

- Microsoft Windows 10
- MATLAB 9.10 R2021a
- Python 3.9 (64-bit)

**Codon Adaptation Index Formula:**

The Codon Adaptation Index (CAI) can be utilized to measure codon bias. The following CAI measure is performed in this study given a set of highly expressed genes.

$$CAI = \left(\prod_{i=1}^{L} w_i\right)^{\frac{1}{L}}$$

where $L$ is the number of codons in the sequence and $w$ is calculated by:

$$w_i = \frac{RSCU_i}{RSCU_{max}}$$

where $RSCU_{max}$ is the highest codon usage frequency for a synonymous codon in a set of highly expressed genes and where $RSCU_i$ is the codon usage frequency for synonymous codon $i$.

**Codon Frequency Distribution Definition:**

The Codon Frequency Distribution (CFD) is calculated using GenScript's Rare Codon Analysis tool accessible via https://www.genscript.com/tools/rare-codon-analysis. This tool has a CFD field which indicates the percentage of codons in a gene sequence that are found to be low-frequency in the host organism (used less than 30% of the time). Thus, we quantify CFD in this study as a percentage.

**Negative Repeat Elements Definition:**

The Negative Repeat Elements are calculated using GenScript's Rare Codon Analysis tool accessible via https://www.genscript.com/tools/rare-codon-analysis. This tool has a field referring to Negative Repeat Elements which are sections of repeating codons that negatively impact expression. Decreasing the number of negative repeat elements will yield a more optimized sequence.

**Negative Cis-Regulatory Elements Definition:**

The Negative Cis-Regulatory Elements are calculated using GenScript's Rare Codon Analysis tool accessible via https://www.genscript.com/tools/rare-codon-analysis. This tool has a field referring to

Negative CIS Elements which are cis-acting elements or motifs in the sequence that negatively regulate gene expression in transcription and translation.
