## Supplementary material for "ICOR: Improving codon optimization with recurrent neural networks": S2 Table

| **Amino Acid** | **Code** | **Integer** |
| --- | --- | --- |
| Alanine | A | 1 |
| Arginine | R | 2 |
| Asparagine | N | 3 |
| Aspartic acid | D | 4 |
| Cysteine | C | 5 |
| Glutamine | Q | 6 |
| Glutamic acid | E | 7 |
| Glycine | G | 8 |
| Histidine | H | 9 |
| Isoleucine | I | 10 |
| Leucine | L | 11 |
| Lysine | K | 12 |
| Methionine | M | 13 |
| Phenylalanine | F | 14 |
| Proline | P | 15 |
| Serine | S | 16 |
| Threonine | T | 17 |
| Tryptophan | W | 18 |
| Tyrosine | Y | 19 |
| Valine | V | 20 |
| Asparagine or Aspartic acid | B | 21 |
| Glutamine or Glutamic acid | Z | 22 |
| Unknown amino acid | X | 23 |
| Translation stop | * | 24 |
| Gap of indeterminate length | - | 25 |
| Unknown character | ? | 26 |
