## Supplementary material for "ICOR: Improving codon optimization with recurrent neural networks": S2 Appendix

**Training Data Specifications**

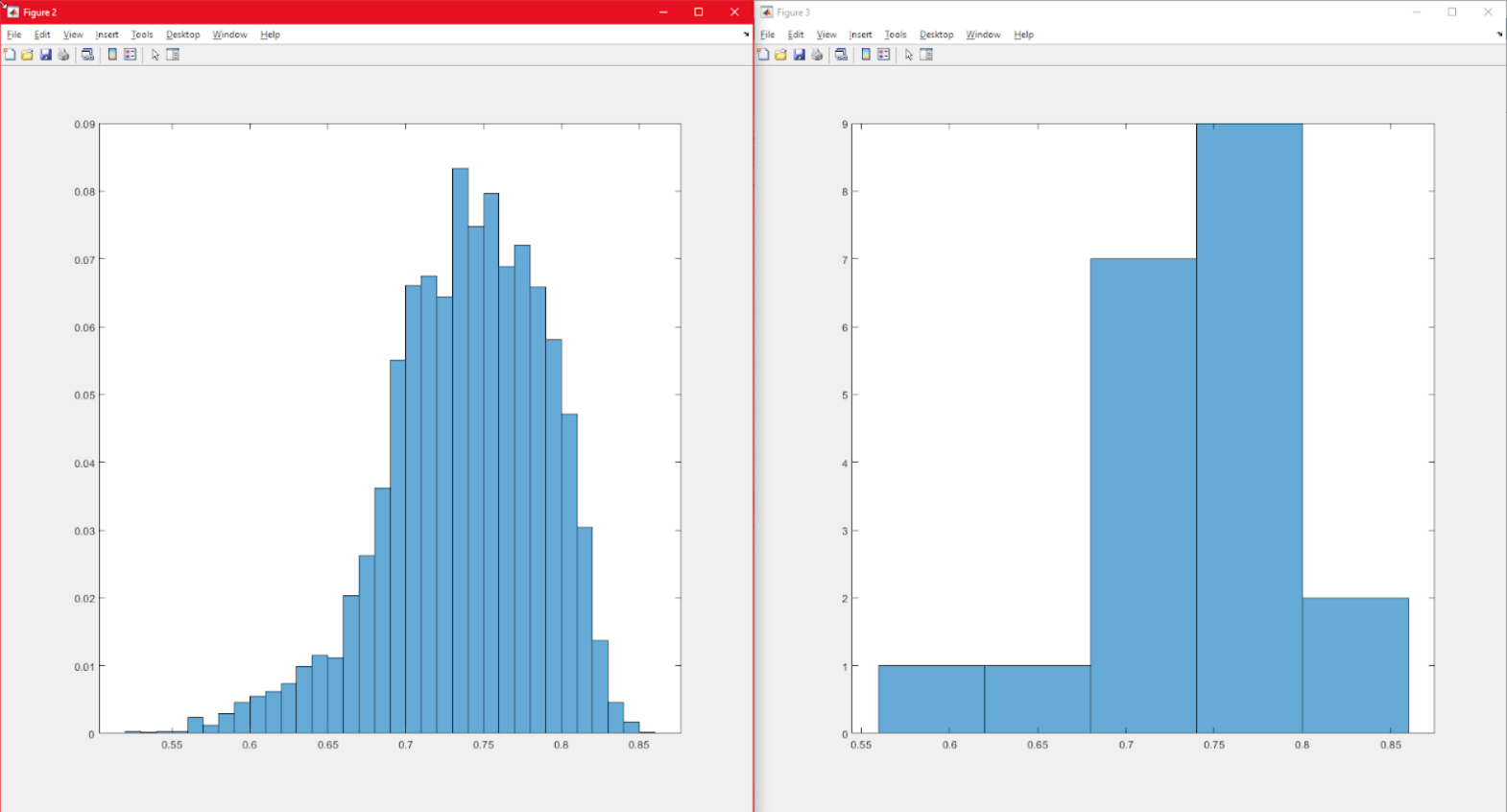

This histogram displays the distribution of the codon adaptation index of the 42,266 genes utilized in the ICOR training dataset. Codon adaptation index is depicted on the x-axis and proportion of overall dataset is depicted on the y-axis.

Of these, the 7,406 genes with the highest codon adaptation index were selected to use as part of the training dataset. We intentionally choose to train only on *E. coli* genes because they will be stable within the chassis during expression. Thus, using these genes to inform codon selection in a synthetic gene may bring about this stability. Because the model only predicts based on genes from *E. coli*, based on evolutionary-instilled processes, these genes naturally expressed will theoretically not be insoluble. However, these genes vary in expression levels.
