## Supplementary material for "ICOR: Improving codon optimization with recurrent neural networks": S3 Table

| **Hyperparameter Used in Model Training** | **Value of Parameter** |
|---|---|
| Gradient Decay Factor (dictates the decay rate of gradient moving average) | 0.95 |
| Learning Rate (the extent to which weights are updated during training, also known as step size) | $5 \times 10^{-5}$ |
| L2 Regularization (factor of weight decay) | 1e-8 |
| Shuffling (shuffles the order of training data) | never |
| Dropout (the factor by which neurons are temporarily ignored during training) | 0.2 |
| Padding (ensures a ragged input—sequences are not padded to a uniform length) | off |
| Sorting (sort input by ascending sequence length) | on |
| Optimizer (the solver algorithm used when training the network) | adam (adaptive moment estimation) |
